# Pharmacological inhibition of the heart of glass (HEG1)-Krev interaction trapped protein 1 (KRIT1) protein complex increases Krüppel-like Factors 4 and 2 (KLF4/2) expression in endothelial cells

**DOI:** 10.1101/744821

**Authors:** Miguel Alejandro Lopez-Ramirez, Mark K. Haynes, Preston Hale, Killian Oukoloff, Matthew Bautista, Brendan Gongol, John Y. Shyy, Carlo Ballatore, Larry A. Sklar, Alexandre R. Gingras

**Affiliations:** Department of Medicine, University of California San Diego, 9500 Gilman Drive, La Jolla, CA, 92093, USA; Department of Pharmacology, University of California San Diego, 9500 Gilman Drive, La Jolla, CA, 92093, USA; Department of Pathology, Center for Molecular Discovery, University of New Mexico School of Medicine, Albuquerque, NM 87131 USA; Skaggs School of Pharmacy and Pharmaceutical Sciences, University of California San Diego, 9500 Gilman Drive, La Jolla, CA, 92093, USA

**Keywords:** HEG1, KRIT1, KLF2, KLF4

## Abstract

The Krüppel-like Factors 4 and 2 (KLF4/2) are transcription factors and master regulators of endothelial cells (ECs) phenotype and homeostasis. KLF4/2 are important blood-flow-responsive genes within ECs that differentially regulate the expression of factors that confer anti-inflammatory, antithrombotic, and antiproliferative effects in ECs. Genetic inactivation of endothelial KRIT1 (Krev interaction trapped protein 1) or HEG1 (Heart of glass) lead to upregulation of KLF4/2 expression levels. Furthermore, increased expression of thrombomodulin (THBD) and suppression of thrombospondin (THBS1) was ascribed to elevation of KLF4/2 as a result of loss of endothelial KRIT1. Here, we developed a high-throughput screening assay to identify inhibitors of the HEG1-KRIT1 interaction and identified, HEG1-KRIT1 inhibitor 1 (HKi1), as a promising hit inhibitor. The crystal structure of HKi1 bound to the KRIT1 FERM domain confirmed the primary screening results and ultimately led to the identification of a fragment-like inhibitor (HKi3), which occupies the HEG1 pocket producing comparable activity. These findings suggest that these inhibitors block the interaction by competing with the HEG1 for binding to KRIT1 FERM domain. Moreover, our results demonstrate that HKi3 upregulates KLF4/2 gene expression in two types of human ECs. These results reveal that acute pharmacological inhibition of the HEG1-KRIT1 interaction rapidly induces expression of KLF4/2 and their important transcriptional targets thrombomodulin and thrombospondin.

## INTRODUCTION

Endothelial cells (ECs) line the entire circulatory system and EC dysfunction plays a central role in the development of vascular disease states such as atherosclerosis and thrombosis. HEG1 is a transmembrane receptor that is required for cardiovascular development in both zebrafish and mammals (1–3). Our previous studies showed that the cytoplasmic domain (tail) of HEG1 binds directly to KRIT1 (also known as CCM1), the protein product of the *KRIT1* gene (4,5). We solved the structure of the complex of KRIT1 FERM domain with HEG1, and used that structure to show that the interaction recruits KRIT1 to cell-cell junctions thereby anchoring the complex to support heart development in zebrafish (4). Both HEG1 and KRIT1 dampen gene expression levels of transcriptional regulators *KLF4/2* (6,7), and therefore play roles in controlling the sensitivity of ECs to hemodynamic forces (8,9). KLF4/2 are strongly activated within regions of pulsatile shear stress (PS) (10). In turn, KLF4/2 can increase expression of genes that encode anticoagulants (e.g. *THBD* encoding thrombomodulin, TM)(11) or vasodilators (e.g. *NOS3* encoding endothelial nitric oxide synthase, eNOS)(12), and suppress expression of genes that antagonize angiogenesis (e.g. *THBS1* encoding thrombospondin1, TSP1) and NFkB-driven proinflammatory genes (e.g. vascular adhesion molecules including, *VCAM1* and *ICAM1)(13).* Because the HEG1 binding pocket on the KRIT1 FERM domain is both discrete and unique, we reasoned that it may be suitable for small molecule drug/probe discovery efforts to identify inhibitors of this protein-protein interaction. Moreover, we hypothesized that pharmacologic inhibition of the HEG1-KRIT1 protein complex could elevate expression of KLF4/2 and their transcriptional targets. Towards this end, we conducted a screen for small molecule inhibitors of the KRIT1-HEG1 interaction, and identified an inhibitor that, when tested in two different EC lines in culture, upregulated KLF4/2, which in turn can upregulate thrombomulin and suprress thrombospondin expression.

## RESULTS

### A new flow cytometry assay to study HEG1-KRIT1 protein interaction

We previously solved the crystal structure of the KRIT1 FERM domain bound to the C-terminal region of the HEG1 cytoplasmic tail (Fig. 1A) (4). Because the HEG1 binding pocket on the KRIT1 FERM domain is both discrete and unique, we hypothesized that specific inhibitors of the HEG1-KRIT1 protein complex could be identified. To test this hypothesis, we developed a high-throughput flow cytometry-screening assay to screen for compounds that block the HEG1-KRIT1 protein-protein interaction. We had previously shown that the HEG1 cytoplasmic tail can be used as an affinity matrix for KRIT1 binding (4) and we also used this matrix to identify important interactors for HEG1 function such as Rasip1 (14). Using a similar approach, we coupled the biotinylated HEG1 cytoplasmic tail (a.a. 1274-1381) peptide to 6-micron SPHERO Neutravidin coated particles (Fig. 1B). We first added varying amounts of biotinylated HEG1 peptide to the beads (Fig. 1C) and addition of purified recombinant GFP-KRIT1 FERM domain to the HEG1 matrix beads, without washes, leads to a dose dependent GFP intensity increase by flow cytometry (Fig. 1D). Importantly, we observed many beads forming doublets at a 1,500 nM HEG1 concentration in the light scattering affecting the GFP signal (Fig. 1C). Therefore, we used a concentration of 150 nM biotinylated HEG1 for the assay, which gives the best signal without aggregation of the beads. Secondly, addition of increasing amounts of purified recombinant GFP-KRIT1 FERM domain to the HEG1 matrix beads, without washes, lead to a dose dependent GFP intensity increase by flow cytometry with EC_50_ = 32.4 nM (Fig. 1E, blue line), showing that GFP-KRIT1 binds the HEG1 tail on the beads. Importantly, a KRIT1(L717,721A) mutant with a major reduction in HEG1 affinity(4), showed a dramatic reduction in binding (Fig. 1E, red line), validating this approach and showing specific binding. Therefore, we used a concentration of 70 nM for the assay. Our previous data using Isothermal Titration Calorimetry (ITC) showed a K_D_ = 1.2 μM for the KRIT1 FERM domain binding to a HEG1 peptides in solution (4). Using our HEG1 matrix beads we observed an EC_50_ = 32.4 nM (Fig. 1E). We observed that the measured apparent off-rate is slower than the actual off-rate and we assumed that is because following dissociation, the GFP-KRIT1 can bind to an unoccupied HEG1 tail before diffusing out of the matrix. Importantly, incubation of the GFP-KRIT1 FERM domain with a non-biotinylated HEG1 C-terminus 7-mer peptide block the interaction in a dose dependent manner with IC_50_ = 410 nM (Fig. 1F). These results establish a reliable and quantitative assay to study the HEG1-KRIT1 protein interaction by flow cytometry.

**FIGURE 1.**
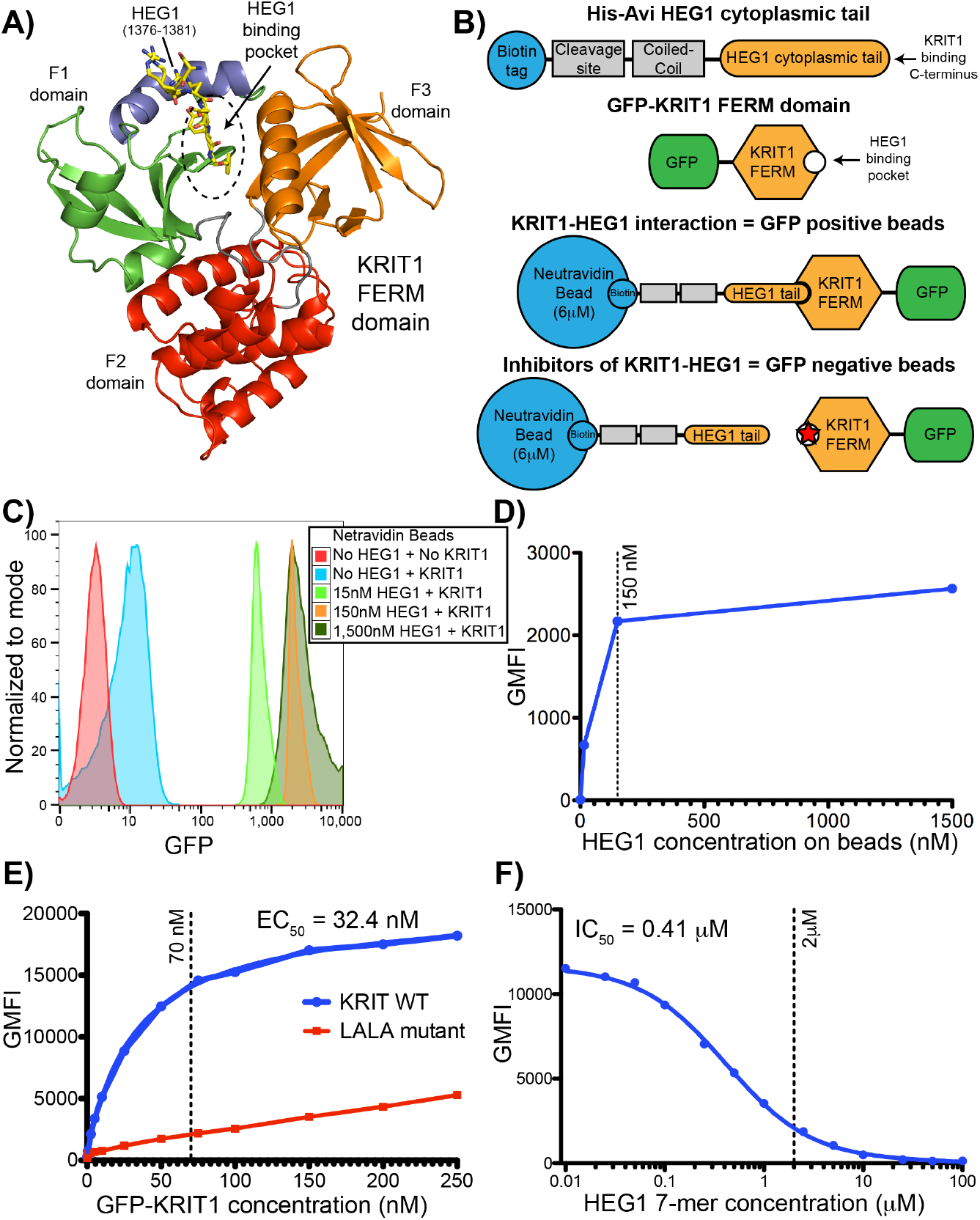
Flow cytometry assay for the HEG1-KRIT1 FERM domain interaction. (A) Ribbon diagram of KRIT1 FERM domain in complex with the HEG1 cytoplasmic tail (PDB ID: 3u7d). The HEG1 peptide is shown in yellow. The KRIT1 FERM domain consists of three subdomains: F1 (green and blue); F2 (red); and F3 (orange). The feature of the F1 domain that is not present in other FERM domain is shown in blue and that region is an important part of the HEG1 binding pocket. (B) Schematic representation of the HEG1 cytoplasmic tail (a.a. 1274-1381) peptide coupled to Neutravidin beads and of the EGFP-KRIT1 FERM domain. (C) Flow cytometry profile of SPHERO Neutravidin Polystyrene Particles coated with increasing amount of biotinylated HEG1 peptide and 150 nM EGFP-KRIT1 FERM domain. We noticed many beads doublets in the light scattering at 1,500 nM concentration of HEG1 peptide. (D) Titration curve for the interaction of EGFP-KRIT1 FERM domain with increasing amounts of HEG1 on the beads as shown in panel C. We used the 150 nM HEG1 peptide concentration for future experiments. (E) Titration curve for the interaction of 150 nM HEG1 on the beads with increasing amounts of EGFP-KRIT1 FERM domain (0-250 nM) wild-type (blue line) and KRIT1(L717,721A) mutant (red line). We used the 70 nM EGFP-KRIT1 concentration for future experiments. (F) Competition binding curve of 70 nM EGFP-KRIT1 FERM domain binding to 150 nM HEG1 on the beads with increasing amounts on non-biotinylated HEG1 7-mer peptide. We used the 2 μM HEG1 7-mer concentration for future experiments.

### High-throughput screening identifies inhibitors of HEG1-KRIT1 protein interaction

Since our flow cytometry assay to study the HEG1-KRIT1 interaction is simple, requires no washes, and can be inhibited using a HEG1 peptide, we scaled it down for high throughput in 384-well plate format. The assay required only 10 μl of sample per well in nanomolar concentrations with a count of 1,000 beads per microliter. We performed a pilot screen using an automated sample loader attached to a flow cytometer and analyzed 2 μl of sample per well (2,000 beads). By alternating beads with GFP-KRIT1 in the absence or presence of 2 μM HEG1 7-mer blocking peptide (Supporting Information, Fig. S1A) we measured a Z’ of 0.528, classifying the assay as excellent (Fig. 2A) (15). Out of 6,026 compounds screened (Supporting Information, Fig. S1B) we identified that HEG1-KRIT1 inhibitor 1 (HKi1) (Fig. 2B) identified as NAD+-dependent deacetylases (16–18), had promising pharmacological properties with an IC_50_ value of ~10 μM (Fig. 2C). However, consistent with a high logP value of 5.7 (Fig. 2B), HKi1 had limited aqueous solubility at 50 μM concentrations or higher in our buffer conditions. As a result, saturating conditions in the assay could not be achieved (Fig. 2C).

**FIGURE 2.**
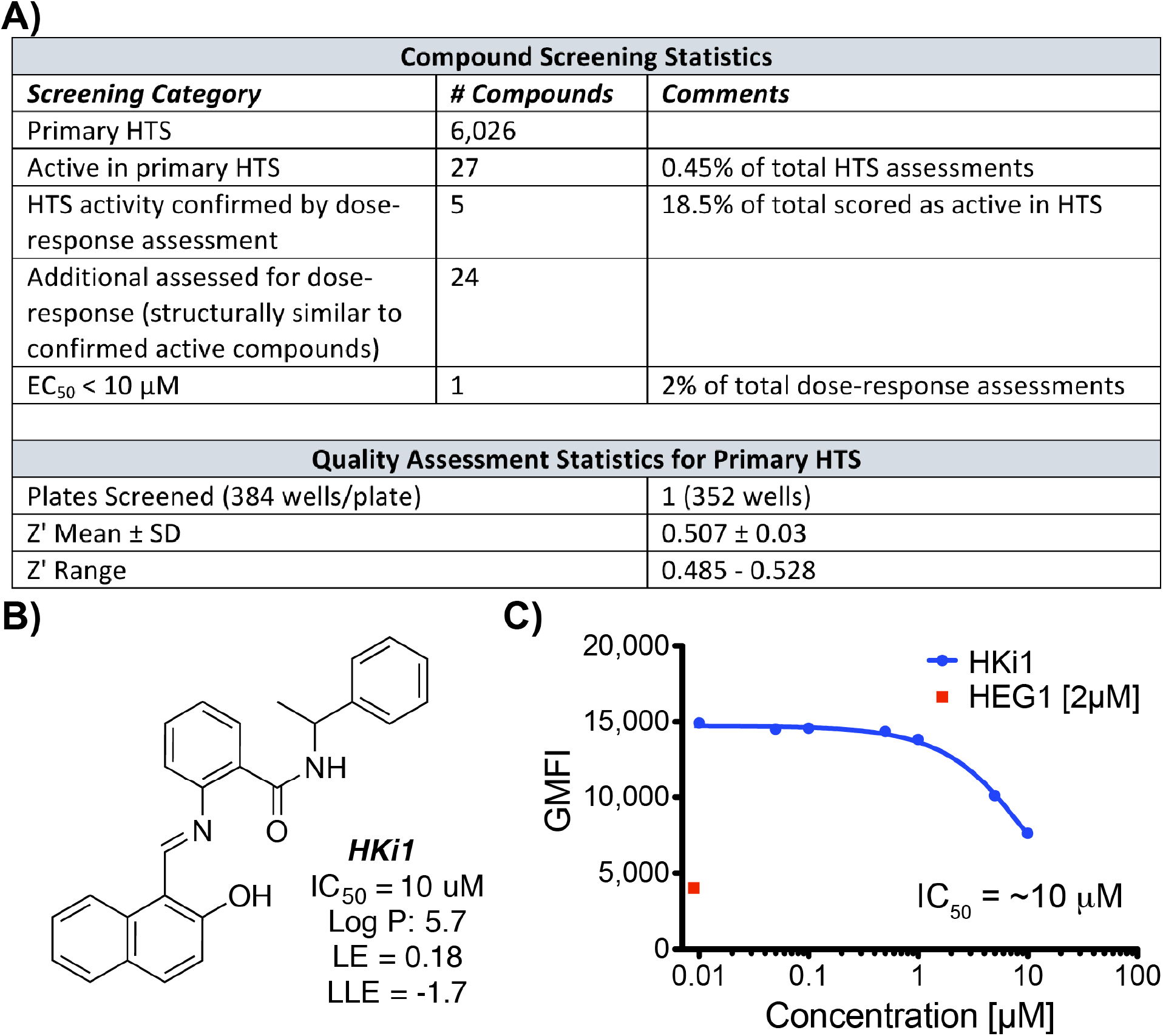
HKi1 is an inhibitor of the HEG1-KRIT1 interaction. (A) Compound screening statistics. (B) Chemical structure of HKi1. LE = (1.37/HA) x pIC_50_ where HA is the number of non H atoms present in the ligand; LLE = pIC_50_ – LogP. (C) Competition binding curve of 70 nM EGFP-KRIT1 FERM domain binding to 150 nM HEG1 on the beads with increasing amounts on HKi1. HKi1 had a poor solubility in our buffer and concentrations >30 μM could not be reached.

### Crystal structure of KRIT1 FERM domain in complex with HKi1

Since we previously determined the crystal structure of the KRIT1 FERM domain bound to a HEG1 peptide (4) (Fig. 1A and 3A), we then crystallized the KRIT1 FERM domain in the presence of HKi1 and solved the structure of the complex to 1.75 Å resolution (Table 1). The structure strongly suggests that HKi1 is a *bona fide* inhibitor and it confirms that this compound occupies the same pocket as the HEG1 (Fig. 3B), supporting that HKi1 blocks the interaction by competing orthosterically with the HEG1 for binding to KRIT1 FERM domain. HKi1 is largely hydrophobic (logP = 5.7), as the HEG1 C-terminal Tyr-Phe residues, and sits in the hydrophobic pocket formed at the interface of the F1 and F3 subdomains of the KRIT1 FERM domain. Interestingly, good electron density was observed for approximately half of the molecule and less well-defined electron density was observed for the other half of the molecule (Fig. 3B and Supporting Information, Fig. S2A), suggesting that modifications to HKi1 could improve binding properties. Moreover, a clear socket is found in the binding pocket (Fig. 3B) and many positively charged residues were found in the proximity of the binding pocket, suggesting that the design of analogs with improved binding affinity as well as improved physicochemical properties may be possible.

**FIGURE 3.**
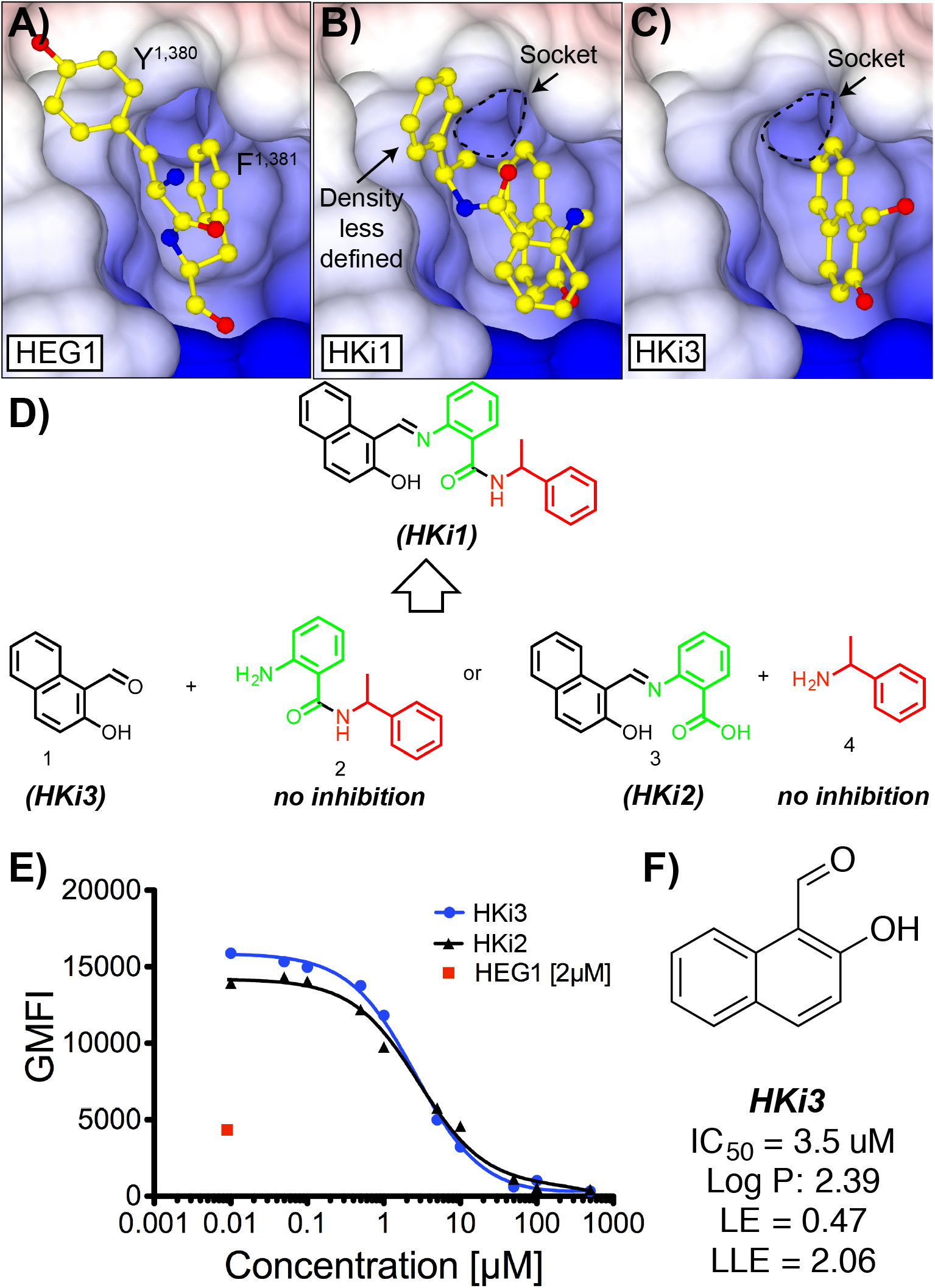
Structure guided HEG1-KRIT1 interaction inhibitors. (A) Surface charge representation of KRIT1 FERM domain in complex with the HEG1 cytoplasmic tail (PDB ID: 3u7d). The HEG1 peptide is shown in yellow with the C-terminal Tyr-Phe sitting in the binding pocket. (B) Crystal structure of the KRIT1 FERM domain in complex with HKi1. The small naphthalene is sitting in the same pocket as the Phe of HEG1 and the electron density for the benzylamine moiety is less defined. (C) Crystal structure of the KRIT1 FERM domain in complex with HKi3. The small naphthalene is sitting in the same pocket as the Phe of HEG1 and good electron density is observed. (D) Chemical structure of HKi1 constituents. (E) Competition binding curve of 70 nM EGFP-KRIT1 FERM domain binding to 150 nM HEG1 on the beads with increasing amounts on HKi2 and HKi3. (F) Chemical structure of HKi3. LE and LLE are described in Fig. 2B. The solubility of HKi3 in aqueous solution is largely improved.

**Table 1.**
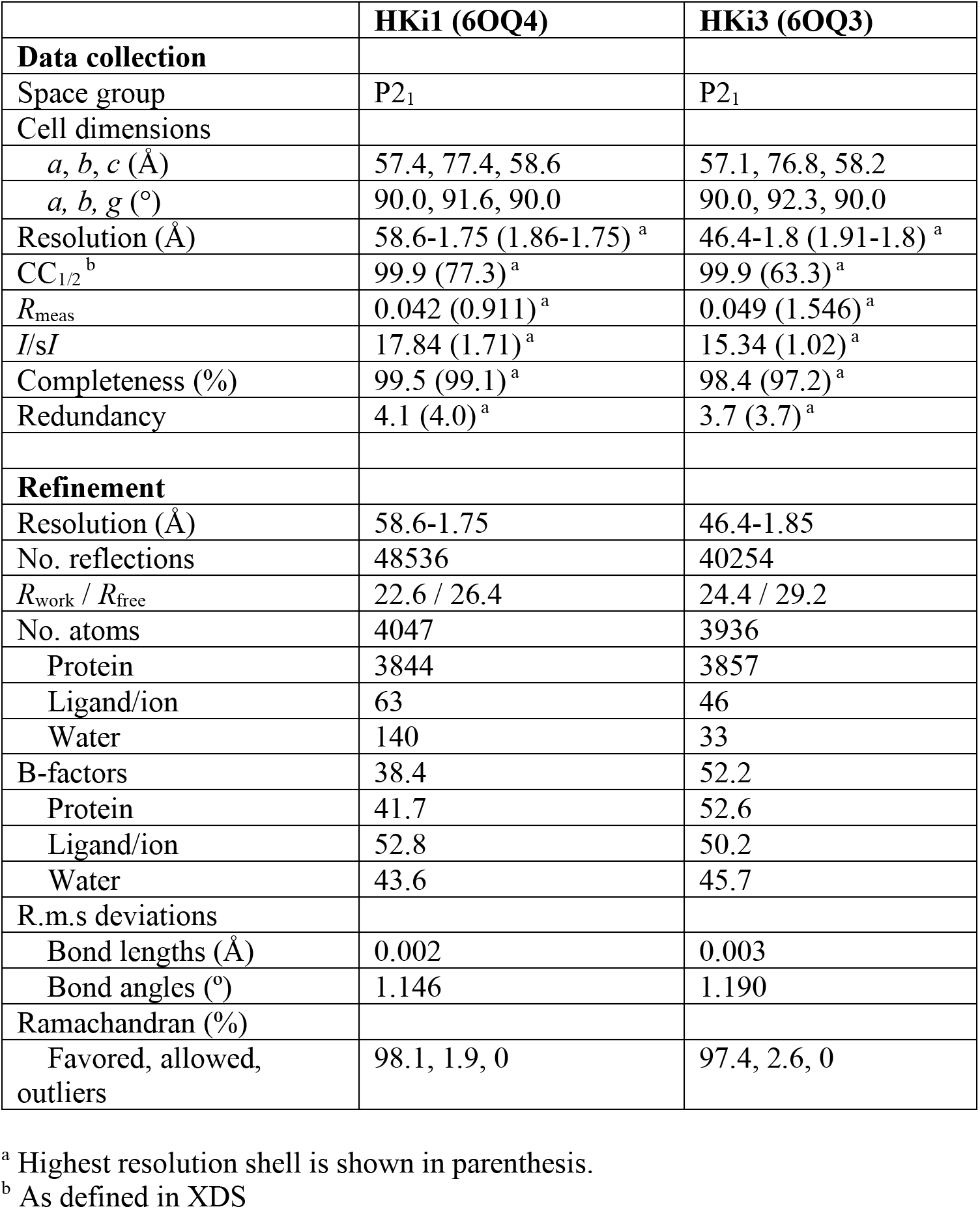
Data collection and refinement statistics for the KRIT1 FERM domain bound to Rap1 and small inhibitors: HKi1 and HKi3.

|  | HKi1 (6OQ4) | HKi3 (6OQ3) |
| --- | --- | --- |
| **Data collection** |  |  |
| Space group | P2<sub>1</sub> | P2<sub>1</sub> |
| Cell dimensions |  |  |
| *a*, *b*, *c* (Å) | 57.4, 77.4, 58.6 | 57.1, 76.8, 58.2 |
| *a*, *b*, *g* (°) | 90.0, 91.6, 90.0 | 90.0, 92.3, 90.0 |
| Resolution (Å) | 58.6-1.75 (1.86-1.75)<sup>a</sup> | 46.4-1.8 (1.91-1.8)<sup>a</sup> |
| CC<sub>1/2</sub><sup>b</sup> | 99.9 (77.3)<sup>a</sup> | 99.9 (63.3)<sup>a</sup> |
| *R*<sub>meas</sub> | 0.042 (0.911)<sup>a</sup> | 0.049 (1.546)<sup>a</sup> |
| *I*/*σI* | 17.84 (1.71)<sup>a</sup> | 15.34 (1.02)<sup>a</sup> |
| Completeness (%) | 99.5 (99.1)<sup>a</sup> | 98.4 (97.2)<sup>a</sup> |
| Redundancy | 4.1 (4.0)<sup>a</sup> | 3.7 (3.7)<sup>a</sup> |
| **Refinement** |  |  |
| Resolution (Å) | 58.6-1.75 | 46.4-1.85 |
| No. reflections | 48536 | 40254 |
| *R*<sub>work</sub> / *R*<sub>free</sub> | 22.6 / 26.4 | 24.4 / 29.2 |
| No. atoms | 4047 | 3936 |
| Protein | 3844 | 3857 |
| Ligand/ion | 63 | 46 |
| Water | 140 | 33 |
| B-factors | 38.4 | 52.2 |
| Protein | 41.7 | 52.6 |
| Ligand/ion | 52.8 | 50.2 |
| Water | 43.6 | 45.7 |
| R.m.s deviations |  |  |
| Bond lengths (Å) | 0.002 | 0.003 |
| Bond angles (°) | 1.146 | 1.190 |
| Ramachandran (%) |  |  |
| Favored, allowed, outliers | 98.1, 1.9, 0 | 97.4, 2.6, 0 |

<sup>a</sup> Highest resolution shell is shown in parenthesis.<sup>b</sup> As defined in XDS**Table 1. Data collection and refinement statistics for the KRIT1 FERM domain bound to Rap1 and small inhibitors: HKi1 and HKi3.**

### HKi3 blocks HEG1-KRIT1 protein-protein interaction

In addition to the relatively high lipophilicity and low aqueous solubility, HKi1 is also characterized by suboptimal values in efficiency metrics, such as the ligand efficiency (LE) and the lipophilic ligand efficiency (LLE) (19,20). These characteristics suggest that this particular compound may be problematic as a starting point for hit-to-lead optimization studies. However, analysis of the complex structure (Fig. 3B) suggested that while the naphthalene moiety of HKi 1 may play an important role in determining the compound’s binding and inhibitory activity, other fragments *(e.g.,* the benzylamine) may not be as intimately involved in the binding to KRIT1. This observation led us to deconstruct HKi1 into its constituent fragments (Fig. 3D) and investigate the ability of these fragments to inhibit the HEG1-KRIT1 *in vitro.* These studies confirmed that substructures containing the substituted naphthalene fragment, such as HKi2 and HKi3, produce inhibition of the HEG1-KRIT1 interaction with IC_50_ values of 3.5 μM that are closely comparable to the IC_50_ value of the parent compound, HKi1 (Fig. 3E). These findings suggested that the 2-hydroxy-1-naphthalene fragment bearing a reactive carbonyl group at the C1 position might be essential for the inhibition of the HEG1-KRIT1 interaction. Interestingly, when we crystallized the KRIT1 FERM domain in complex with HKi3 (Fig. 3C, Table 1 and Supporting Information, Fig. S2B) we observed that the naphthalene fragment retained the same binding mode within the HEG1 binding pocket on KRIT1 as observed in the HKi1 complex (Fig. 3B). In this regard, recent studies found that fragments of ligands that fully overlap with the strongest hot spot generally retain their position and binding mode when the rest of the molecule is removed (21). Although we found no direct evidence of covalent modification of the aldehyde moiety of HKi3, it is conceivable that the chemically reactive aldehyde or imine moiety present in these naphthalene inhibitors may be involved in reversible covalent binding with KRIT1 leading to relatively potent inhibition of the HEG1-KRIT1 interaction. Not surprisingly, given the relatively small size and reduced lipophilicity of HKi3 compared to HKi1 (Fig. 3F), the LE as well as the LLE are considerably improved suggesting that the former compound may be considered as a promising starting point for further optimization.

### HKi3 upregulates KLF4 and KLF2 levels in endothelial cells

Since loss of endothelial KRIT1 increases expression of KLF4 and KLF2 transcription factors (7–9,22,23), we next evaluated whether the small molecule HKi3 affect endothelial KLF4 and KLF2 gene expression. To this end, HUVEC and a human brain endothelial cell-line, hCMEC/D3, were treated with HKi3 and KLF4 and KLF2 expression levels were determined (Fig. 4). We observed that KLF4 and KLF2 mRNA levels were upregulated following addition of 25 μM HKi3 to the culture medium in hCMEC/D3 cells for 12 h of treatment (Fig. 4A and 4B). Increasing the concentration of HKi3 further upregulated KLF2 and KLF4 mRNA levels (Fig. 4A and 4B). While upregulation of KLF4 mRNA levels (~6.5 fold increase at 50 μM) were pronounced compared with controls (Fig. 4B), the changes in KLF2 mRNA levels (~2.3 fold increase at 50 μM) were moderate but significant (Fig. 4A). We also noted that incubation of hCMEC/D3 cells with 50 μM HKi3 induced a rapid upregulation of KLF2 (~ 1.5 fold increase) as early as 4 h and continued to be increased until the end of the experiment at 24 h (Fig. 4C). Moreover, KLF4 mRNA levels (~3.5 fold increase) were also upregulated at 4 h after treatment and higher KLF4 levels (~6 fold increase) were detected following 24 h of treatment (Fig. 4D).

**FIGURE 4.**
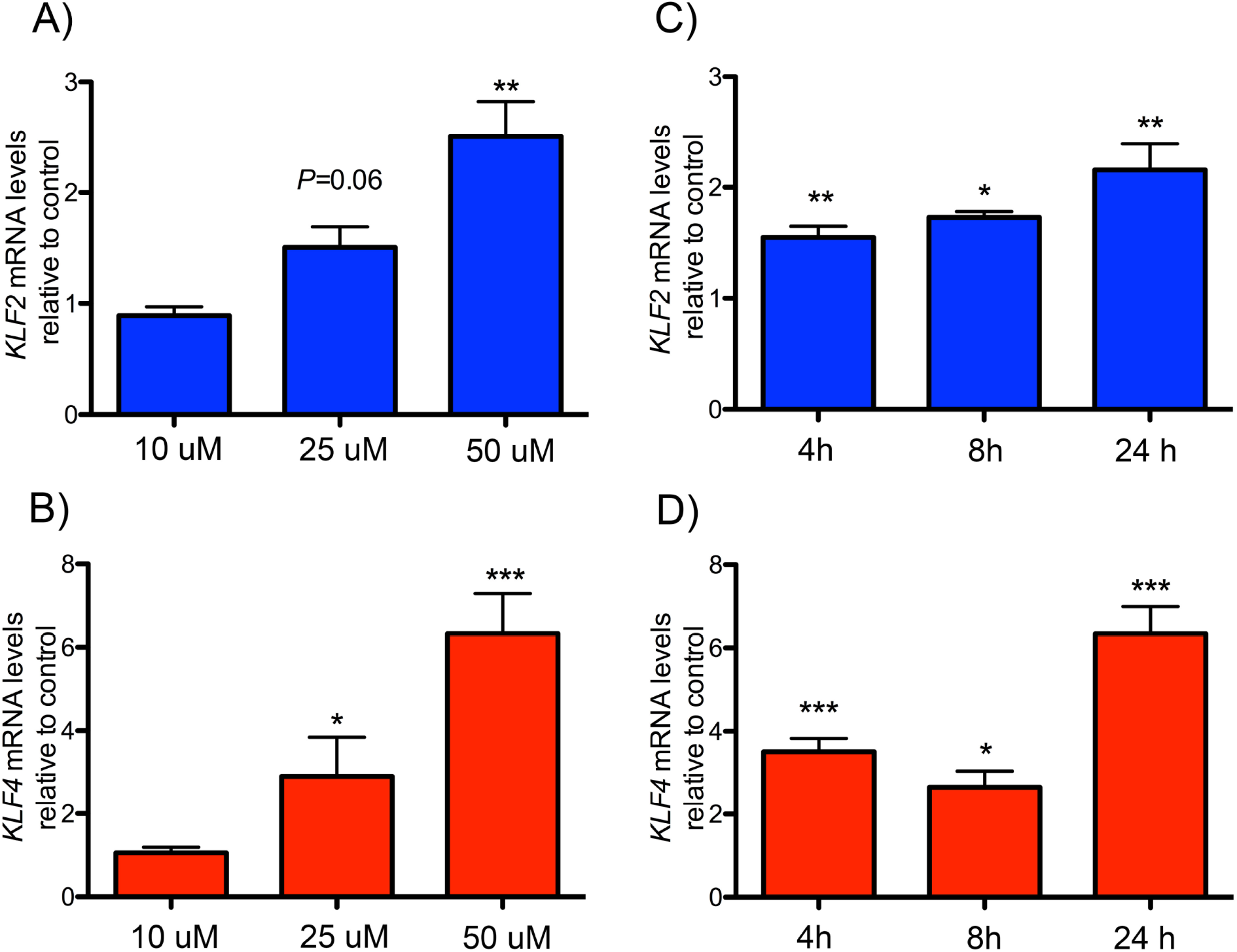
HKi3 treatment leads to KLF2 and KLF4 upregulation in hCMEC/D3 endothelial cells. (A-D) mRNA levels of HKi3 treated cells as determined by qPCR. (A,B) Dose response of (A) KLF2 and (B) KLF4 mRNA expression at indicated doses for 12 hours. HKi3 induces KLF2 and KLF4 mRNA expression at indicated concentrations. (C,D) Timecourse, HKi3 (50 μM) induces a rapid and sustained upregulation of (C) KLF2 and (D) KLF4 mRNA expression. (A-D) Bar graphs represent mRNA levels relative to vehicle control ± SEM (n = 3, 2-tailed *t* test). *, P < 0.05; **, P < 0.01; ***, P < 0.001.

Similar results were observed using HUVEC cells. We observed that incubation of HUVEC with HKi3 upregulated both KLF4 and KLF2 expression in a dose-dependent manner (Fig. 5A and 5B). More importantly, we found after 24 hours incubation with 50 μM HKi3 (Fig. 5C), an 8-fold upregulation of *THBD* and a 6-fold downregulation of *THBS1,* which have been ascribed to elevation of KLF4/2 expression. These results indicate that pharmacological inhibition of KRIT1-HEG1 protein interaction can be used to study the early signaling events that lead to upregulation of KLF4 and KLF2 and their important transcriptional targets such as *THBD* and *THBS1.*

**FIGURE 5.**
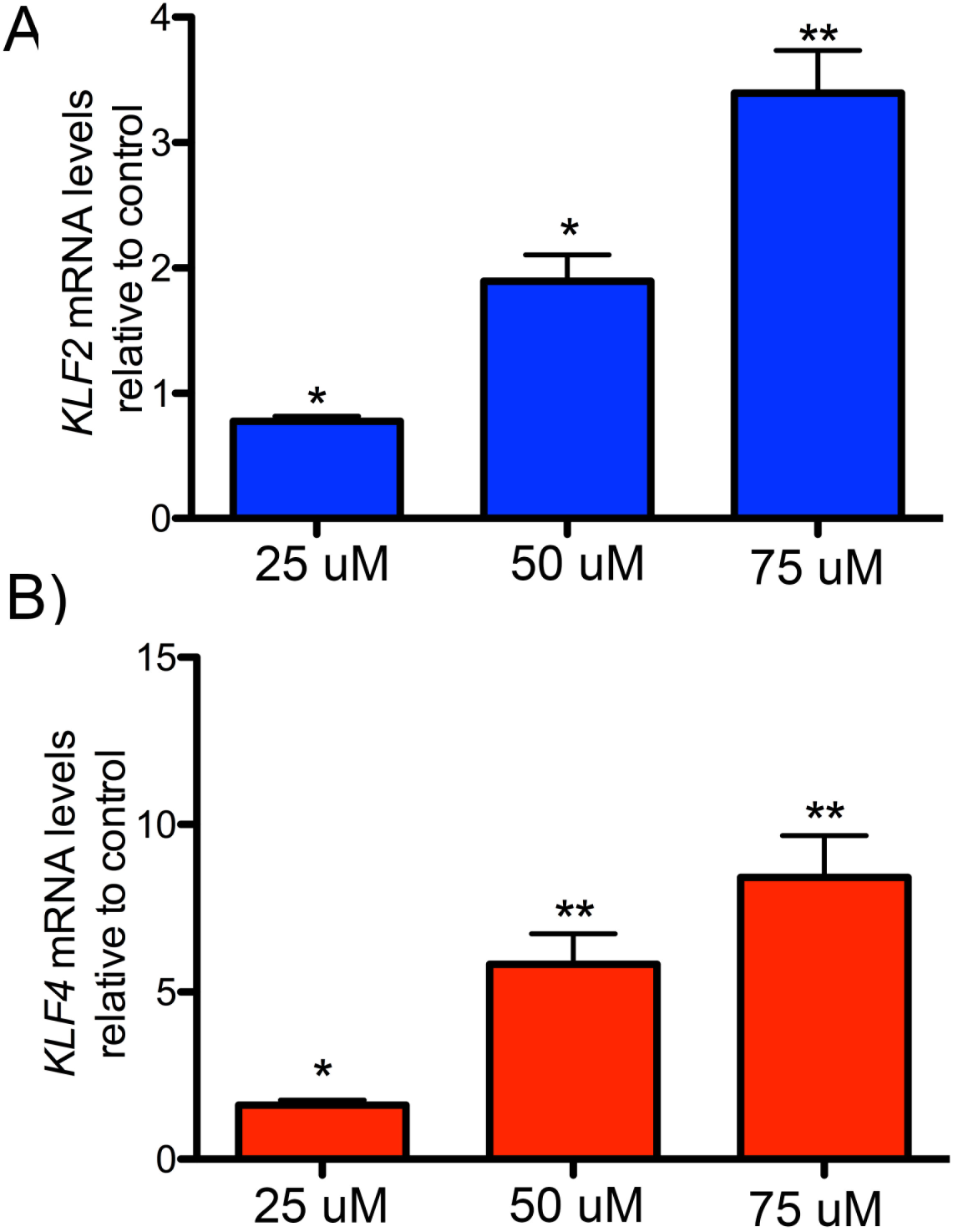
HKi3 treatment leads to KLF2 and KLF4 upregulation in HUVECs. (A-B) mRNA levels of HKi3 treated HUVEC as determined by qPCR. (A,B) Dose response of (A) KLF2 and (B) KLF4 mRNA expression at indicated doses for 24 hours. HKi3 induces KLF2 and KLF4 mRNA expression at indicated concentrations. (A-B) Bar graphs represent mRNA levels relative to vehicle control ± SEM (n = 3, 2-tailed *t* test). *, P < 0.05; **, P < 0.01; ***, P < 0.001.

## DISCUSSION

Many of the effects of pulsatile sheer stress (PS) are due to upregulation of transcription factors KLF4 and KLF2, which in turn can increase expression of genes that encode anticoagulant cofactors (e.g., THBD encoding thrombomodulin, TM) and suppress expression of genes that antagonize angiogenesis (e.g.,THBS1 encoding thrombospondin1, TSP1). Thus, PS can upregulate KLF4/2 in endothelium and their important transcriptional targets (9,24–29).

Loss of KRIT1 leads to cerebral cavernous malformations (CCM) (30). However, there is abundant evidence from murine models that perinatal endothelial-specific inactivation of Krit1 leads to CCM (6,9,31), whereas genetic inactivation of endothelial Krit1 in adults does not. Moreover, HEG1 mutations have never been identified in human CCM and deletion of HEG1 in mice does not cause CCM (1). Here we developed a method for pharmacological inhibition of the HEG1-KRIT1 protein-protein interaction and found the predicted upregulation of *KLF4* and *KLF2* gene expression, and expected effects on some of their target genes such as *THSP1* and *THBD,* as we previously observed with knockdown of KRIT1 in EC(9,29). Moreover, KLF4/2 were upregulated within 4 hr of drug addition, a time long before acute genetic inactivation of *Krit1* would do so. Thus, these compounds will provide a new tool for analysis of signaling pathways downstream of loss of HEG1-KRIT1 interaction and may provide a new pharmacological approach to upregulate KLF4/2.

### Screening methodology

Here we described a high-throughput screening using 6,026 unbiased small molecules. The HEG1 cytoplasmic tail was bound to NeutrAvidin coated polystyrene particles and GFP-KRIT1 FERM domain binding was measured by flow-cytometry. The assay was aimed to uncover inhibitors of the HEG1-KRIT1 interaction; indeed it led to the identification of a small molecule that specifically inhibits this interaction. This simple flow-cytometry assay provides a platform for screening for small molecule modulators of the HEG1-KRIT1 interaction and possibly of other protein-protein interactions by using a similar approach. The screen is high-throughput, required minimum labor, and is easy to score for positive. Furthermore, the screen gave remarkably specific results, as is evident by the fact that the hit was crystallized with the KRIT1 FERM domain.

### Identification of HEG1-KRIT1 inhibitors

An examination of the crystal structure of KRIT1 bound to HKi1 indicated that the substituted naphthalene fragment may play a critical role in establishing interactions with the HEG1 binding pocket on KRIT1. Indeed, deconstruction of HK1 into its primary constituents revealed that smaller 2-hydroxy-naphthalenes bearing a reactive carbonyl moiety such as an imine (HKi2) or an aldehyde (HKi3) can inhibit the HEG1-KRIT1 interaction with comparable IC_50_ values as HKi 1. The low μM IC_50_ values of these smaller fragments, especially HKi3, appears to be considerably more potent (i.e., approximately ~100-1000 times) than those typically observed for low MW fragments that can establish only a few non-covalent interactions with the target protein. This observation suggests that the relatively reactive carbonyl moiety of HKi3 may undergo covalent reversible binding with the KRIT1 FERM domain. Among the 20 proteinic amino acids, the side chains of lysine and arginine are capable of forming covalent reversible interactions with aldehydes (typically in the form of an iminie or enamine adducts). The crystal structure shows that the HEG1-binding pocket of KRIT1 contains three lysine residues (K475, K724, and K720). Therefore, it is conceivable that the aldehyde moiety of HKi3 may engage in covalent reversible binding with one of these residues leading to the relatively potent inhibition of the HEG1-KRIT1 interaction. However, thus far we have been unable to obtain direct evidence of covalent modification.

### HKi3 as a lead compound

An exciting possibility will be to generate specific inhibitors of the HEG1-KRIT1 interaction by modifying the basic structure of our lead compound. The crystal structure of the KRIT1 FERM domain bound to HKi3 shows that the compound occupies the same position as the Phe^1,361^ of HEG1 (Fig. 3A and 3C) and importantly this small fragment molecule retains all the biological activity of HKi1. This suggests that the Phe^1,361^ of HEG1 site is a hot spot that could be exploited for drug discovery. To this end, the structure shows a socket in the HEG1-binding pocket that could be targeted by appropriately growing the 2-hydroxy-1-naphthaldehyde fragment identified in this study (Fig. 3C). Furthermore, the structure of the KRIT1 FERM domain bound to the HEG1 cytoplasmic tail (Fig. 3A) shows where the HEG1 Tyr^1,360^ sits in the pocket and suggests that extending HKi3 with a Tyr-like moiety could significantly improve the compound specificity and affinity. The crystal structure with HKi1 (Fig. 3B) shows that a moiety of the molecule almost reaches this area, but that some restraints in the molecule prevents it to have the right conformation to occupy that place. The three crystal structures of the KRIT1 FERM domain suggest opportunities for modifications of HKi3 to improve its binding. Thus, our results confirm that HKi3 is a *bona fide* inhibitor and suggest that the binding site may be amenable to small molecule drug discovery programs.

In conclusion, we designed a screen, found an inhibitor of the KRIT1-HEG1 interaction, and found that inhibition of this signaling pathway can upregulate the transcription factors KLF4/2 that are very important in vascular integrity. New pharmacological agents to upregulate KLF4/2 have the potential of confer anti-inflammatory properties that are predicted to be beneficial in diseases such as atherosclerosis. This pharmacological manipulation of the HEG1-KRIT1 mainly upregulates KLF4 in contrast to other therapeutic approaches such statins that preferentially upregulates KLF2(32). The combination of the two approaches could complement each others in future therapeutics.

## EXPERIMENTAL PROCEDURES

### Plasmid construction and protein purification

HEG1 intracellular tail model protein was prepared as previously described (4). In brief, His6-tagged HEG1 intracellular tail containing an *in vivo* biotinylation peptide tag at the N-terminus was cloned into pET15b, expressed in BL21 Star (DE3) and purified by nickel-affinity chromatography under denaturing conditions. Synthetic human non-biotinylated HEG1 7-mer peptide (residues 1375–1381) was purchased from GenScript. His6-EGFP-KRIT1(WT) FERM domain (417–736) and KRIT1(L717,721A) mutant were cloned into pETM-11 and expressed in BL21 Star (DE3). Recombinant His-EGFP-KRIT1 was purified by nickel-affinity chromatography, and further purified by Superdex-75 (26/600) size-exclusion chromatography (GE Healthcare). The protein concentration was assessed using the A280 extinction coefficient of 71,740 M^-1^.

Human KRIT1 FERM domain, residues 417–736 was expressed and purified as described previously (5). Briefly, KRIT1 was cloned into the expression vector pLEICS-07 (Protex, Leicester, UK) and expressed in *Escherichia coli* BL21 Star (DE3) (Invitrogen). Recombinant His-tagged KRIT1 was purified by nickel-affinity chromatography; the His tag was removed by cleavage with tobacco etch virus protease overnight, and the protein was further purified by Superdex-75 (26/600) size-exclusion chromatography. The protein concentration was assessed using the *A*280 extinction coefficient of 45,090 M^-1^.

Human Rap1 isoform Rap1b (residues 1–167) cloned into pTAC vector in the *E. coli* strain CK600K was the generous gift of Professor Alfred Wittinghofer (Max Planck Institute of Molecular Physiology, Germany). The Rap1b was expressed and purified as described previously (33). The protein concentration was assessed using a molar absorption coefficient of A280 = 19,480 M^-1^ as previously reported (34).

Equimolar concentrations of KRIT1 FERM domain and GMP-PNP loaded Rap1b were mixed and loaded on a Superdex-75 (26/600). The column was pre-equilibrated and run with 20 mM Tris, 50 mM NaCl, 3 mM MgCl2, and 2 mM DTT (pH 8). The final complex concentration was determined using a molar absorption coefficient of A280 = 61,310 M^-1^ for the KRIT1-Rap1b complex.

### Bead Coupling

SPHERO Neutravidin Polystyrene Particles, 6-8 μM (Spherotech) were washed twice with wash buffer (20 mM Tris, 150 mM NaCl, pH 7.4 containing 0.01% NP-40, and 1mM EDTA). Prior to incubation with biotin-tagged HEG 1 cytoplasmic tail protein, an appropriate volume of bead slurry was passivated to inhibit non-specific binding by incubation for 30 minutes at room temperature in reaction buffer containing 0.1% BSA (20 mM Tris, 150 mM NaCl, pH 7.4 0.01% NP-40, 1mM EDTA, 1mM DTT, and 0.1% BSA). Passivated beads were collected by centrifugation, resuspended to 3,600 particles/μl in reaction buffer and biotinylated HEG1 tail was added to a final concentration of 150 nM and incubated overnight on a rotator at 4°C. The beads were washed three times by centrifugation with ice-cold reaction buffer to remove unbound HEG1 peptide. Beads were diluted such that a final concentration of 2000 beads/μL were available for addition to assay plates.

### Flow cytometry assay

A final volume of 100 μl containing 140 nM EGFP-KRIT1 FERM domain, with 10% DMSO or 10% compounds in DMSO, was incubated for 15 minutes at room temperature on a rotator. 100 μl of beads were added to the mixture for a final volume of 200 μl at 1,000 particles/μl with 70 nM EGFP-KRIT1 and incubated for 15 minutes at room temperature on a rotator. The control beads were: without KRIT1 (minimum signal); with KRIT1 (maximum signal); and with KRIT1 plus 2 μM HEG1 7-mer (positive blocking control). The EGFP fluorescence was measured using a BD Accuri flow cytometer. For screening purposes, the final volume of the reaction was scaled down to 10 μl and samples were processed as previously described (35).

### Assay Plate Assembly

Plate assays were performed in 384-well microtiter plates (Greiner Bio-one, #784101). Reaction buffer, HEG1-coupled beads, and EGFP-KRIT-FERM constructs were added using a MultiFlo^TM^ Microplate Dispenser (BioTek Instruments, Inc.). Compounds were added to single-point assay plates pre-loaded with reaction buffer using a Biomek^NX^ liquid handler (BeckmanCoulter) equipped with a 100 nL pintool (V & P Scientific, Inc.). Compound libraries were dispensed to a final concentration of 10 μM. An equal volume (10 nL) of DMSO was added to the vehicle control wells. Following the addition of library compounds, 5 μL of assay buffer was added and the plates were mixed before addition of 5 μL of the protein-coupled bead mixtures; Plates were protected from light and incubated on a rotator for 15 minutes at room temperature. Binding of EGFP-KRIT to HEG1 coupled beads was evaluated using an Accuri C6 flow cytometer.

Dose response plates were assembled similarly. In this instance, test compounds were added to dose response plates using a dilution protocol of the acoustic dispenser that resulted in a final concentration range of 100 – 0.015 μM.

### Data Acquisition

Assay plates were sampled using the HyperCyt^TM^ high throughput flow cytometry platform (Intellicyt; Albuquerque, NM). During sampling, the probe moves from well to well and samples 1 – 2 μL from each well pausing 0.4 sec in the air before sampling the next well. The resulting sample stream consisting of 384 separated samples is delivered to an Accuri C6 flow cytometer (BD Biosciences; San Jose, CA). Plate data are acquired as time-resolved files that are parsed by software-based well identification algorithms and merged with compound library files. Plate performance was validated using the Z-prime calculation (15). Compounds that satisfied hit selection criteria in the primary screen were cherry-picked from compound storage plates and tested to confirm activity and determine potency. Dose response data points were fitted by Prism software (GraphPad Software Inc., San Diego, CA) using nonlinear least-squares regression in a sigmoidal dose-response model with variable slope, also known as the 4-parameter logistic equation. Curve fit statistics were used to determine the concentration of test compound that resulted in 50% of the maximal effect (EC_50_), the confidence interval of the EC_50_ estimate, the Hill slope, and the curve fit correlation coefficient.

### Crystallization of the KRIT1-Rap1b-HKis complexes

The purified KRIT1 FERM domain-Rap1b complex at 8.25 mg/ml was used for crystallization. Crystals were grown at room temperature using the sitting-drop method by mixing equal volumes of protein complex and reservoir solution (2 + 2 μl). The reservoir solution contained 20-25% PEG 3,350, 100 mM Tris, pH 8.5, 100 mM KCl. After 1 week or later, ~0.5 μl of 10 mM compounds in DMSO was added to the drop for 1 day. The crystals were briefly transferred to the reservoir solution containing 20% glycerol before freezing in liquid nitrogen.

### Structure Determination

Diffraction data for the KRIT1 FERM domain-Rap1b-HKis complexes were collected at the Advanced Light Source beamline 5.0.3. The data were processed with XDS (36). The structures were solved by molecular replacement using Phaser with the structure of the KRIT1-Rap1b complex (PDB ID: 4hdo).

### Cell culture

hCMEC/D3 cells at passages 30–37 were grown to confluence on collagen-coated plates and cultured using in EGM-2 MV medium and supplemented with complements obtained from the manufacturer (Lonza) as previously reported (37). HUVEC (Lonza) at passages 4-7 were grown to confluence on gelatin-coated plates and maintained using complete EGM-2 media (Lonza). HKi002 was maintained at room temperature for 30 min rotating before use. Cells were then treated with HKi002 at the concentrations and times indicated for each experiment. Cells were maintained at 37 C in 95% air and 5% CO2.

### Reagents

All reagents were from Sigma (St Louis, MO) unless otherwise indicated. Plastic-ware was from VWR (Radnor, PA) and Greiner Bio-One (Monroe, NC). Neutravidin Bead sets for were from Spherotech, Inc., (Lake Forest, IL). All solutions were prepared with ultra-pure 18 MΩ water or anhydrous DMSO. Flow cytometric calibration beads were from Bangs Laboratories Inc., (Fishers, IN) and Spherotech, Inc. Off patent commercial libraries were purchased from Prestwick Chemical (Illkirch-Graffenstaden, France), SelleckChem (Houston, TX), Spectrum Chemical (New Brunswick, NJ), and Tocris Bio-Science (Bristol, UK). We also purchased a collection of on patent drugs from MedChem Express (Monmouth Junction, NJ) that was specifically assembled by UNM collaborators. All purchased libraries were provided as 10 mM stock solutions in 96-well matrix plates except the MedChem Express library which was provided as individual powders that were subsequently solubilized in DMSO. All libraries were reformatted using a Biomek FX^P^ laboratory automated workstation into 384-well plates for storage (Greiner #784201; Labcyte #PP-0200). Low volume dispensing plates (Labcyte #LP-0200) were assembled using an Agilent BioCell work station (Santa Clara, CA). Low volume dispensing plates (Labcyte #LP-0200) were assembled using an Agilent BioCell work station (Santa Clara, CA). The following compounds were purchased from: HKi1: Sirtinol (Selleckchem); 2-hydroxy-1-naphthaldehyde (Ark Pharm); and 2-amino-N-(1-phenylethyl)benzamide (Enamine).

## ACKNOWLEDGMENTS

This research used resources of the Advanced Light Source, which is a DOE Office of Science User Facility under contract no. DE-AC02-05CH11231. This work was supported by the National Institutes of Health grants K01HL 133530 (to M.A.L.-R.), and NS 092521 (to M.H.G.), the UCSD Academic Senate RS164R-GINGRAS (to A.R.G.), and UC Multi-campus Research Program MRP-17-454909 (to A.R.G. and C.B.). Thanks to Mark H. Ginsberg for valuable discussions. Data deposition: The atomic coordinates and structure factors have been deposited in the Protein Data Bank, www.wwpdb.org (PDB ID code 6OQ4, and 6OQ3).

