## Supplemental for "Pharmacological inhibition of the heart of glass (HEG1)-Krev interaction trapped protein 1 (KRIT1) protein complex increases Krüppel-like Factors 4 and 2 (KLF4/2) expression in endothelial cells"

Supplementary Information for

### Pharmacological inhibition of the Cerebral Cavernous Malformation protein complex increases KLF2 and KLF4 in endothelial cells.

Miguel Alejandro Lopez-Ramirez, Mark K. Haynes, Preston Hale, Killian Oukoloff, Matthew Bautista, Carlo Ballatore, Larry A. Sklar, and Alexandre R. Gingras

Alexandre R. Gingras


**This PDF file includes:**

Supplementary text
Figs. S1 to S2
Tables S1
References for SI reference citations

#### Supplementary Information Text

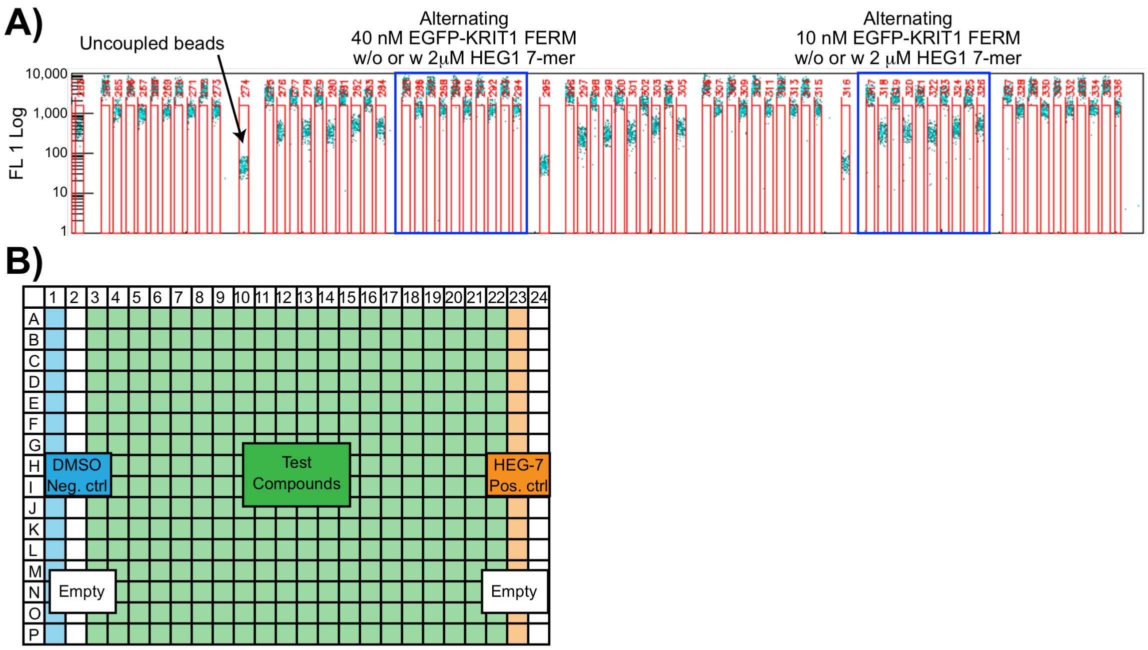


FIGURE S1. **Hyper-Cyt high throughput flow cytometry.** (A) Samples from wells are delivered to the flow cytometer (red boxes) as a series of 2 µL volumes separated by air gaps. Beads in each sample generate a pulse of events (~2000 beads/sample) as they pass through the point of detection in the flow cytometer. Gaps between peaks represent the passage of air bubbles. Each peak receives a time stamp corresponding to its time of appearance (horizontal axis). Boundaries are determined for each sample peak (red lines), and well IDs (horizontal axis labels) are assigned on the basis of time stamps. Green fluorescence intensity (FL) of individual beads (dots) is plotted for each well. (B) Cartoon illustrating the plate layout. The compounds were used at a final concentration of 10 µM and the HEG1 7-mer peptide at 2 µM.

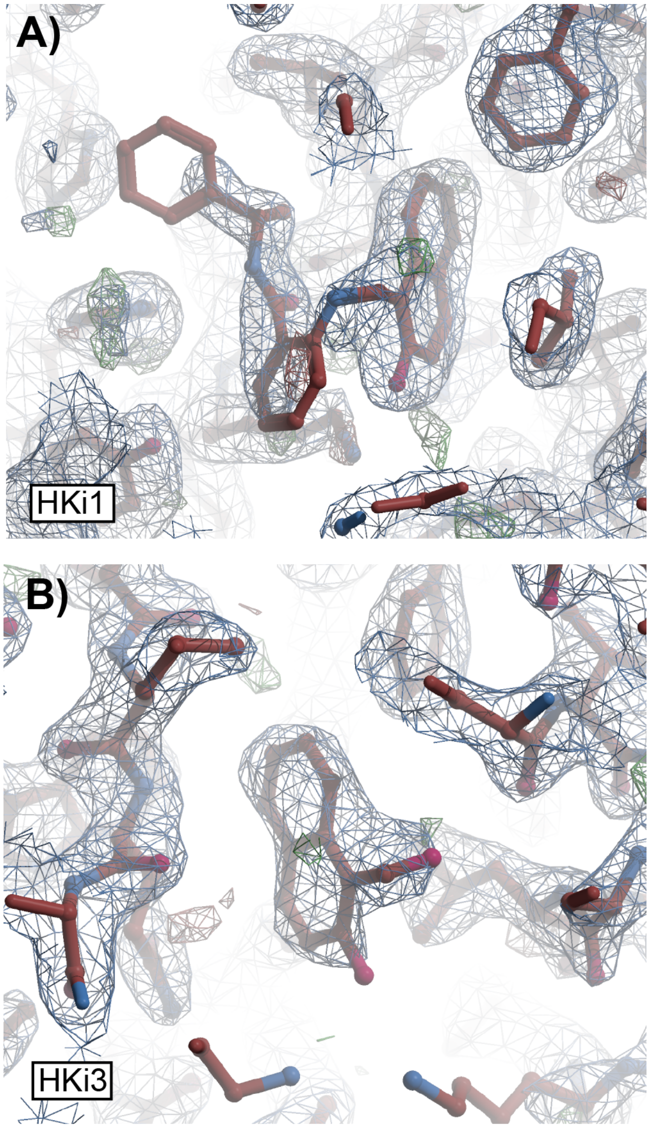


FIGURE S2. **Structural characteristics of the KRIT1 complexes.** (A-B) the KRIT1 FERM domain bound to: A) HKi1; and B) HKI3. Refined 2F0-FC map (blue) and F0-FC at 1σ and 3 σ respectively (red and green) show a good match between the structure and the electron density.

**SI References**
